# Evaluating geographic variation within molecular operational taxonomic units (OTUs) using network analyses in Scandinavian lakes

**DOI:** 10.1101/2020.08.06.240267

**Authors:** Dominik Forster, Guillaume Lentendu, Monica Wilson, Frédéric Mahé, Florian Leese, Tom Andersen, Maryia Khomich, Micah Dunthorn

## Abstract

Operational taxonomic units (OTUs) are usually treated as if they are internally uniform in environmental metabarcoding studies of microbial and macrobial eukaryotes, even when the OTUs are being used to infer biogeographic patterns. The OTUs constructed by the program Swarm have underlying network topologies in which nodes represent amplicons and edges represent 1 nucleotide differences between nodes. Such networks can be exploited to search for biogeographic patterns within each OTU. To do this, here we used an available protistan metabarcoding dataset consisting of the hypervariable V4 region of the 18S rRNA locus amplified from 77 lakes collected across Norway and Sweden. The 82 most abundant and wide-spread OTUs constructed by Swarm were evaluated using shortest path, assortativity, and geographical analyses. We found that while pairs of amplicons from the same lake were usually connected directly to each other within the OTUs, these pairs of amplicons from the same lake did not form assortative clusters within the OTUs, and amplicons were not more connected with other amplicons occurring in neighboring lakes than expected by chance. This new approach to looking at within-OTU is applicable to other metabarcoding datasets and we provide code to perform these analyses.

## INTRODUCTION

Molecular environmental sequencing methodologies are used to evaluate the biodiversity and biogeography of microbial and macrobial eukaryotes. One of these methods is metabarcoding, where specific gene regions are primer-amplified then sequenced (Deiner et al., 2017; Taberlet, Bonin, Zinger, & Coissac, 2018). Metabarcoding studies found, for example, that protists in soils and sediments can have extensive biogeographic patterns (D. Forster et al., 2015; Lentendu et al., 2018; Ritter et al., 2019; Venter, Nitsche, Domonell, Heger, & Arndt, 2017), global terrestrial fungal richness is decoupled from plant diversity (Tedersoo et al., 2014), and marine nematodes and protists can adhere to radically different macroecological processes (Fonseca et al., 2014). In metabarcoding studies the sequencing reads are clustered into operational taxonomic units (OTUs), which can be thought as representing species and which are used as the units of comparison in downstream biodiversity and biogeographical analyses (Bik et al., 2012; Santoferrara et al., 2020).

In these biodiversity and biogeography analyses, OTUs are usually treated as if they are internally uniform. However, since Mayr (1942) taxonomists have known that widespread species are not homogeneous, and that intraspecific variation can be geographically structured between different populations. It is this intraspecific variation that may lead to speciation (Coyne & Orr, 2004). And it is this within-OTU variation that can be evaluated to increase the taxonomic resolution in environmental sequencing studies if we are to uncover the biodiversity and biogeography within widely distributed eukaryotic species. Methodologies have therefore begun to be developed to evaluate within-OTU variation using standard population genetic approaches (Elbrecht, Vamos, Steinke, & Leese, 2018; Turon, Antich, Palacín, Præbel, & Wangensteen, 2019).

One approach for evaluating within-OTU variation is to use network analyses. Network analyses allow biologists to perform fine-grained analyses of similarities between reads, because they exploit the information provided by the topology of connections between reads (Bapteste et al., 2013; D. Forster et al., 2015; Watson et al., 2019). Sequence similarity networks are inclusive graphs that provide a unified comparative framework for reads that can be analyzed with methods and tools of graph theory (Junker & Schreiber, 2008; Watson et al., 2019). In such graphs, the nodes represent the objects to be compared (such as reads that have been dereplicated into strictly identical amplicons), with each pair being linked by an edge if there is significant similarity between the two corresponding nodes (e.g., a minimum sequence similarity). These networks provide a first structuring of the data by partitioning it, since the continuity and discontinuity of resemblances between sequences as guided by chosen parameters generally produces distinct subgraphs, called connected components. Connected components display a variety of mathematical and topological properties that can be exploited in comparative investigations such as in: shortest path analyses, where the minimum distance in the network between nodes that share the same attribute (e.g., geographic origin) is recorded; assortativity analyses, where the tendency of nodes that share the same attribute to be connected with each other in the network is assessed; and what we are here calling geographic analyses, where the distance of the geographic origin of nodes connected in the network is compared.

Such network analyses can easily be applied to OTUs that were constructed by the program Swarm (Frédéric Mahé, Rognes, Quince, de Vargas, & Dunthorn, 2014, 2015). Rather than using a global clustering threshold, Swarm uses a local clustering threshold with an iterative single-linkage approach to construct OTUs. Resulting OTUs therefore do not have a pre-determined radius of genetic similarity (i.e., they are not clustered at global similarity values such as 97% or 98%). OTUs constructed by Swarm have underlying network topologies that can be exported using the -i and -j options. Forster et al. (2016) exploited these network topologies underlying Swarm OTUs, by developing an approach to discover novel genetic diversity in already observed OTUs when sampling new environments. The same study also suggested that these networks’ underlying Swarm OTUs can be exploited to look at within-OTU variation in a geographical context. Underlying network topologies of Swarm OTUs structured by geographic variation may indicate allopatric speciation processes. Since, within such geographically structured Swarm OTUs, amplicons if the same geographic origin would be genetically more similar than amplicons of different geographic origin.

One dataset that can be used to demonstrate Forster et al.’s (2016) suggested approach of evaluating within-OTU variation in a geographical context, is the planktonic protistan metabarcoding data from Khomich et al. (2017). In their study, Khomich et al. (2017) originally sampled 77 homogeneous, low-productive boreal lakes across southern Norway and Sweden. DNA isolated from lake epilimnions were amplified with general eukaryotic primers amplifying the hypervariable-V4 region of the 18S-rRNA locus. These metabarcoding data showed that a longitudinal gradient explained more of the differences in composition and diversity of the protist communities than environmental factors, which supported the earlier microscopic studies of Hessen et al. (2006) and Ptacnik et al. (2010).

Here we demonstrate a method to evaluate geographic variation within Swarm OTUs using network analyses (Figure 1). If dispersal is low and isolation high throughout evolutionary time scales, in a Swarm OTU that is structured by geographic variation, amplicons (which are reads de-replicated at 100% similarity *sensu* Mahé et al. (2015)) originating from the same geographic region should be more similar, and therefore preferentially linked to each other rather than to amplicons originating from a different geographic region; consequently, most amplicons in that Swarm OTU will have a neighboring amplicon from the same geographic region and the average shortest path distance to link two amplicons originating from the same geographic region should be small. We used Khomich et al.’s (2017) southern Norway and Sweden lake metabarcoding data, and after re-clustering with Swarm, we analyzed the 82 most abundant and widespread OTUs. We then investigated if there are underlying geographic variation within those OTUs by asking three questions. First, using shortest path analyses, if amplicons from the same lake are connected directly to each other within the OTUs. Second, using assortativity analyses, if amplicons from the same lake form assortative clusters within the OTUs. Third, using geographic analyses, if amplicons are connected with other amplicons occurring in neighboring lakes.

**FIGURE 1.**
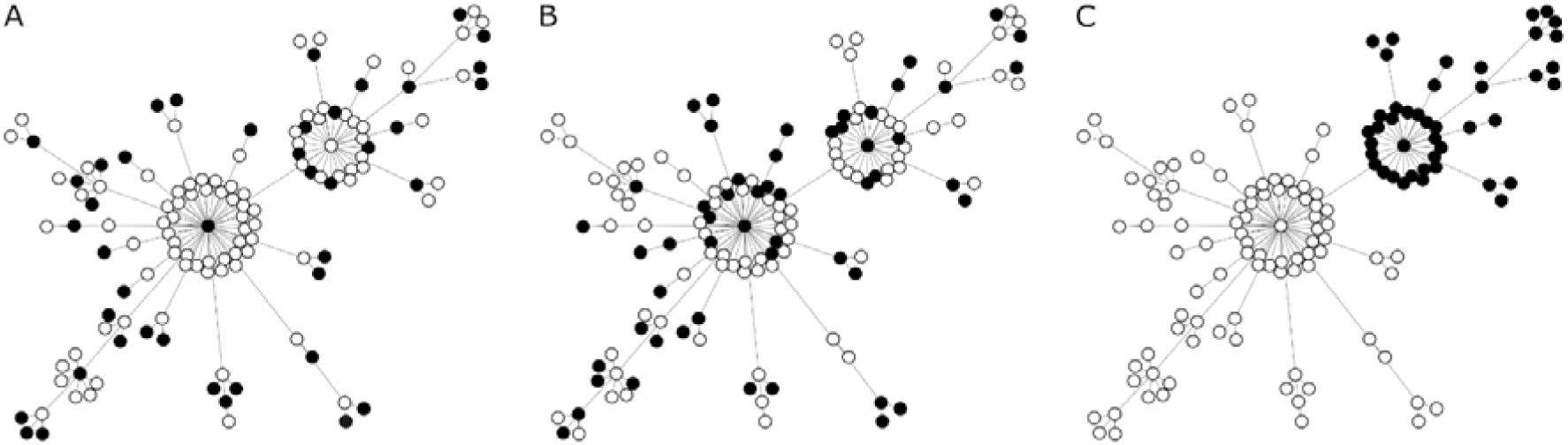
Exemplary illustration of assortativity patterns in a Swarm OTU. Panel A shows a disassortative mock distribution of amplicons within OTU #70 (highlighted in black). The assortativity of the 63 highlighted amplicons in the disassortative mock network scenario was -0.4831. Panel B shows the observed distribution of amplicons from Lake Halsjøen within OTU #70 (highlighted in black). The OTU consisted of 133 amplicons, of which 55 occurred in lake Halsjøen. Their assortativity within this OTU was -0.1604. Panel C shows an assortative mock distribution of amplicons within OTU #70 (highlighted in black), when assuming an underlying biogeographic structuring of the OTU. The assortativity of the 43 highlighted amplicons in the assortative mock network scenario was 0.9826.

## MATERIALS AND METHODS

### Data acquisition

Epilimnic water samples were collected from selected lakes as previously described (Khomich et al., 2017). In brief, for each lake, 15 L of surface water (0-5 m) was filtered through a 100 µm filter and cells were accumulated on nine filters of 2 µm mesh size. Seventy-seven lakes were sampled and analyzed including: Gjersjøen, Halsjøen, Klämmingen, Milsjön, Näsrämmen, Sæbufjorden, Skattungen, Storsjön, Torrsjøn, Vatsvatnet, Vinjevatn, Visten, Vostervatnet (Figure S1). DNA was extracted from one to two filters of each lake and the V4 hypervariable region of the 18S rRNA was amplified with general eukaryotic primers (Stoeck et al., 2010). Raw sequencing data were obtained by 454 Roche pyrosequencing (GS FLX Titanium) and corresponding mapping files were accessed via Dryad (doi:10.5061/dryad.7s6s8).

### Clustering and OTU choice

Raw reads were fully reprocessed through an automatic bioinformatic pipeline for HPC (DeltaMP v0.4, https://github.com/lentendu/DeltaMP). The full bioinformatic procedure can be reproduced using the provided configuration file (File S1). Raw reads were demultiplexed by allowing a maximum of 2 mismatches on the 12 nucleotides long barcode sequences using QIIMEv1.9.1 (Caporaso et al., 2010). Individual .sff files were converted into fasta and quality files. Primers were detected and removed from both ends using Cutadapt v1.18 (Martin, 2011) with an overlap of at least 2/3 of the primer sequence length and allowing up to 6 mismatches on the primer sequences. Only reads containing both primers were kept and reads starting with the reverse primer were reverse-complemented. Reads were filtered using MOTHUR v1.43.0 (Schloss et al., 2009). Reads shorter than 300 nucleotides and longer than 550 nucleotides, with any ambiguous base calls, with homopolymers longer than 8 nucleotides, and with an average Phred score below 20 were discarded. All high-quality filtered reads were dereplicated across the entire dataset to create amplicons, sorted by decreasing abundance, then clustered into operational taxonomic units (OTUs) using the fastidious option *-f* and *-d* 1 in Swarm v3.0.0 (Frédéric Mahé et al., 2014, 2015). Additionally, Swarm’s *-i* and *-j* options were enabled to export the internal structure and the network of each OTU. The seed sequence of each Swarm OTU was used for taxonomic assignment against the PR2 reference database (Guillou et al., 2013) using VSEARCH v2.13.6 (Rognes, Flouri, Nichols, Quince, & Mahé, 2016). Chimeras were detected using VSEARCH and subsequently removed. The full OTU table (Dataset S1), the OTUs internal structures (Dataset S2), the OTUs network file (Dataset S3) and the dereplicated amplicon count table (Dataset S4) are available in the supplements.

All downstream analyses were conducted in R v3.5.1 (R Core Team, 2018; File S1). Ten of the lakes were sequenced more than once; these technical replicates were pooled by lake. OTUs occurring as singletons or doubletons on the global level were removed from downstream analysis (19.9% of all reads; 95.5% of all OTUs). Basic diversity analyses were conducted using the Vegan package (Oksanen et al., 2016). Amplicons assigned to non-protist hits were excluded; e.g. Streptophyta, Metazoa, Fungi, and unidentified Opisthokonta. A subset of the data containing 82 of the most abundant (reads ≥ 0.2% of the total read count in each sample they occurred) and most widespread OTUs (occurring in ≥ 33% of all samples) was finally selected for statistical network analysis.

### Network analyses

For each widespread protist OTU, an undirected network was constructed using Swarm’s internal structure file (i.e. output of option *-i*), which outputs the minimal spanning tree of each OTU network structure as an input for the R package Igraph v1.2.2 (Csardi & Nepusz, 2006). To retrieve the full spanning tree of the internal OTU network structure, additional edges linking two amplicons with one nucleotide difference were retrieved from Swarm’s network file (i.e. output of option *-j*) and added to the network. The grafted edges added by the Swarm fastidious option and corresponding to two nucleotide mismatches were removed in order to keep consistent networks with one nucleotide difference edges only. In this final network, each amplicon is equivalent to one node and each single-linkage between two amplicons is equivalent to one edge representing a single nucleotide difference between the amplicons. Since each OTU represents one enclosed group of amplicons, they are equivalent to connected components within the network (Watson et al., 2019). We defined OTUs with underlying biogeographic structure as connected components in which amplicons from the same lake or from a geographically nearby lake are preferentially connected to each other. And we defined OTUs without underlying biogeographic structure as connected components in which amplicons are randomly connected to each other. By comparing the network structure of each Swarm OTU against randomized network structures in which edges were randomly placed while keeping the node degrees (i.e. the number of edges for each node), we tested if the observed network structures were significantly different from a random structure.

We evaluated the network topology within each OTU using three different concepts from graph theory. First, we evaluated the OTU networks using shortest path analyses. Shortest path analyses measure the distance between any two nodes in the network, expressed as the minimal number of edges that have to be crossed to connect two nodes (Newman, 2003). Neighboring nodes that are directly connected in the network have a shortest path distance of 1. Two nodes that are connected via a third node have a shortest path distance of 2. And nodes in different connected components of the network have a shortest path of ‘infinite distance’ because a path can never be established. For each of the 77 lakes under study, we recorded the minimal shortest path distance of each amplicon to the closest other amplicon from the same lake within the network (avoiding self-loops). In OTUs with underlying biogeographic structure most if not all amplicons from one lake should be neighbors and therefore have a shortest path distance of 1 to the closest other amplicon from the same lake; by contrast, in OTUs without underlying biogeographic structure most amplicons from the same lake will have shortest path distances greater than 1 to each other (including infinite path distances). For amplicons of each lake, we tested if the observed mean shortest path distance fell within a 95% confidence interval of mean shortest path distances obtained from 1,000 randomized networks containing the same number of nodes and the same node degree.

Second, we evaluated the OTU networks using assortativity analyses. Assortativity measures the degree to which nodes of the same type connect with each other in a network, with values ranging from 1 to -1 (Newman, 2003). An example of different assortativity patterns is illustrated in Figure 1. Values close to 1 describe a pattern in which nodes of the same type preferentially connect with each other (Figure 1C), whereas values close to -1 describe a pattern in which nodes of different types preferentially connect with each other (Figure 1A). Values close to 0 indicate a random mixing of node types without preferential connection patterns (Figure 1B). For each of the 77 lakes we ran assortativity analyses, in which the amplicons were categorized into two types: occurring in that particular lake, or not occurring in that particular lake. OTUs with underlying biogeographic structure would yield high assortativity values for at least one lake, because amplicons from that particular lake would be preferentially connected to each other in the OTUs. OTUs without underlying biogeographic structure would yield assortativity values close to 0 or smaller for all lakes under study, because amplicons from different lakes would be randomly connected to each other in the OTUs. We conducted for each lake one assortativity value for the complete network (across all OTUs). This value reflects a general trend observed from all OTUs in the samples. However, since biogeographic patterns in the same habitats may be observed for one OTU but not for another, we also conducted for each lake one separate assortativity analysis within each of the 82 OTUs. Within each Swarm OTU (and for the complete network of all Swarm OTUs), we tested if the observed assortativity for amplicons of the same lake fell within a 95% confidence interval of assortativity values obtained from 1,000 randomized networks containing the same number of nodes and the same node degree.

Third, the assortativity analyses were further conducted in a framework including geographic distances between lakes. In order to take the occurrence of a single amplicon in multiple lakes into account, the centroid geographic position of each amplicon was calculated as the geographic centroid of all lakes in which an amplicon occurs using the package Sp v1.3-1 (Bivand, Pebesma, & Gómez-Rubio, 2013) and Geosphere v1.5-10. In OTUs with underlying biogeographic structure the geographic distance between geographic centroids of neighbor amplicons would be relatively short; by contrast, in OTUs without underlying biogeographic structure the geographic distance between geographic centroids of neighbor amplicons would be random. To test this hypothesis, we compare centroid geographic distances of neighboring amplicons in the observed and randomized networks. The geographic centroid position of each amplicon remained identical in observed and random networks, but the geographic distances between neighbor nodes in the randomized networks were recomputed after shuffling the edges. The set of observed geographic distances between neighbor amplicons in each OTU were compared to each of the 1,000 random networks using a Mann-Whitney test. P-values were corrected for multiple comparisons using the Benjamini & Yekutieli method (Benjamini & Yekutieli, 2001). Significant biogeographic structure was determined when the centroid geographic distances of neighbor amplicons were significantly shorter than expected by chance for 95% or more of the bootstraps in a null model (i.e. p ≤ 0.05).

## RESULTS

### Widespread OTUs

The bioinformatic approach used here produced a similar protist OTU richness (312 ± 107; Figure S1) and rarefaction results (Figure S2) as originally published (Khomich et al., 2017). Among the 2 669 protist OTUs (240 646 reads), the 82 most abundant and widespread OTUs covered 67.9% of total reads (163 382). The widespread OTUs represented on average 20% of each lake’s OTU richness (± 4.1 sd; Figure S1). The taxonomy of the widespread OTUs reflects the total taxonomic composition of the dataset, with the most abundant groups being Cryptophyta, Dinoflagellata, Ochrophyta and Ciliophora (Figure S3).

### Shortest path analyses

Shortest path analyses were performed separately for each lake across all OTUs to record the shortest path distance between two amplicons from the same lake (Table S1).The minimal shortest path distance of 1 between amplicons of the same lake was most often observed for Skattungen (n=1041) and least often for Storsjön (n=223). The maximal shortest path distance between amplicons from the same lake was recorded at a distance of 10 for amplicons from Visten. On average, amplicons from lake Vostervatnet were located closest to each other in the OTUs (average shortest path distance of 1.12), while amplicons from Vatsvatnet were located furthest from each other in the OTUs (average shortest path distance of 1.63). For all lakes, the observed shortest path distance between amplicons of the same lake was shorter than the 95% confidence interval obtained from 1,000 randomized shortest path distances.

In terms of the relative numbers for all lakes, most of the shortest paths with a distance of 1 were observed for Vostervatnet (87.4% of the shortest paths between the amplicons of this lake) and the least shortest paths with a distance of 1 were observed for lake Vatsvatnet (53.3% of the shortest paths between the amplicons of this lake). For all lakes, the majority of amplicons was directly connected with each other in the OTUs and thus differed by a single nucleotide. However, we also observed 855 cases in which only a single amplicon from one lake was in an OTU with amplicons from other lakes. This finding is equivalent to the shortest paths with an infinite distance summarized among all lakes. Most of these cases were registered for Klämmingen (20 times). Regarding relative numbers, the most shortest paths with an infinite distance were observed for lake Gjersjøen (4.6% of the shortest paths between the amplicons of this lake) and the least shortest paths with an infinite distance were observed for Skattungen (0.1% of the shortest paths between the amplicons of this lake).

### Assortativity analyses

For the complete OTU network, the assortativity of amplicons from the same lake ranged from 0.0513 for Vostervatnet to -0.1425 for Torrsjøn (Figure 2). Neither indications for assortative grouping of nodes (values close to 1), nor for disassortative grouping (values close to -1) were observed. Instead, the range of assortativity values indicates a random grouping of nodes (mean value of -0.0744, median of -0.0842) and no preference of amplicons from the same lake to be connected with each other. Nevertheless, the observed assortativity for each lake was still significantly different compared to assortativity values obtained from 1,000 randomized structures of the same network. Out of the 77 assortativity values obtained from the network, the observed assortativity was in 70 cases lower than the 95% confidence interval of the randomized assortativity values; in 7 cases the observed assortativity was higher than the 95% confidence interval of the randomized assortativity values.

**FIGURE 2.**
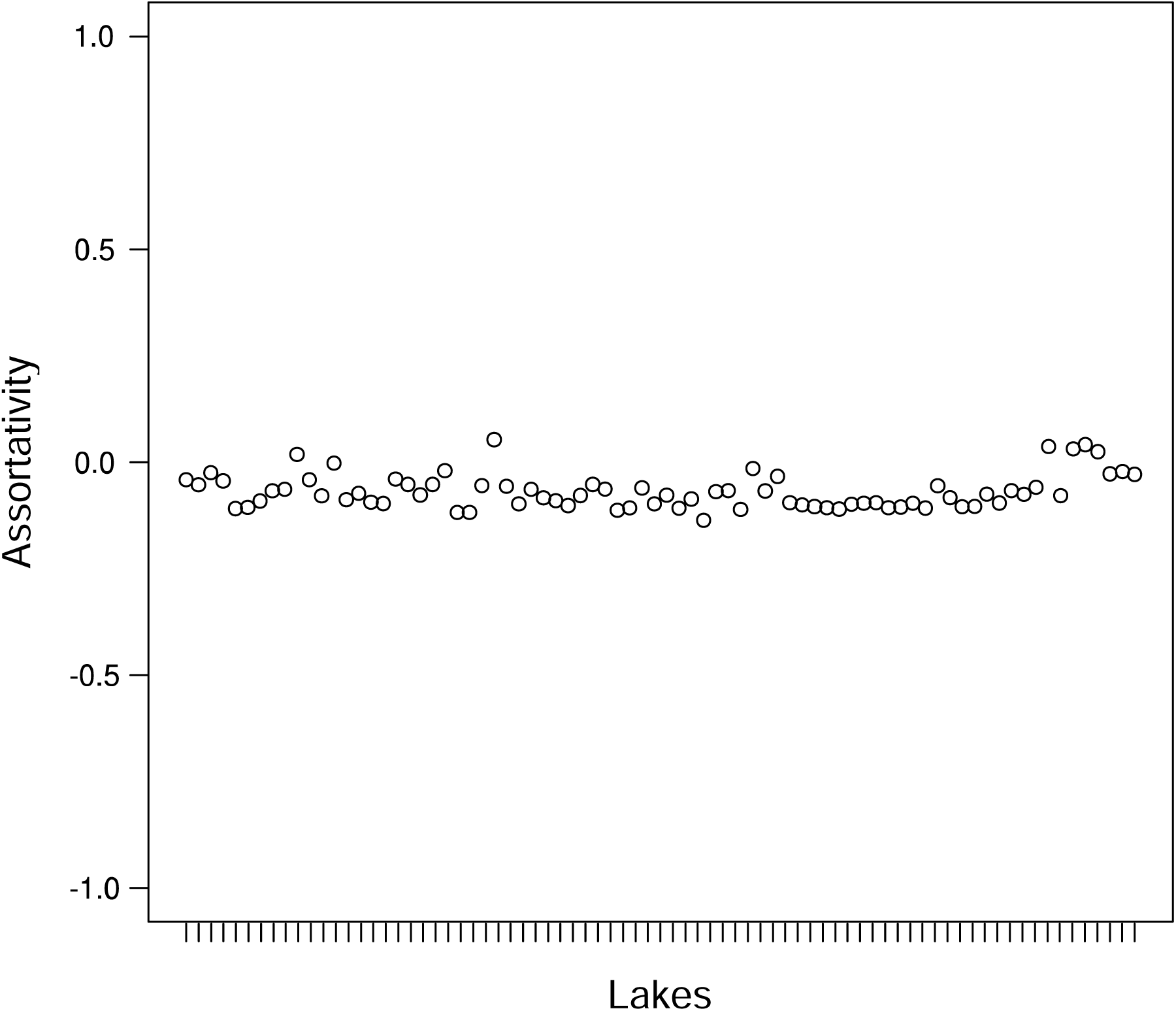
Assortativity of amplicons in the complete OTU network. Calculated independently for each lake across all OTUs (i.e. one assortativity value of each lake for all OTUs).

Regarding assortativity values calculated for individual OTUs, 95% of the values fell within a range of 0.0241 and -0.3561, with a mean value of -0.1644 and a median of -0.1761 (Figure 3). These values further indicate a random grouping of nodes, with a slight tendency towards disassortative grouping. In individual OTUs, amplicons from the same lake were more often than not connected to amplicons from a different lake, rather than to each other. Figure 1B illustrates this case exemplary for the distribution of amplicons from Halsjøen in OTU #70, reaching an assortativity of -0.1604. Nevertheless, when few amplicons from the same lake occurred within one OTU, it was likely that high or low assortativity values were observed, indicating assortative or disassortative grouping for these small groups of amplicons (Figure 4). For example, the highest observed value was at 0.6638 in OTU #48 and the lowest value was at -0.4316 in OTU #86 (Figures 3 and 4). In the first case, this value reflects the connection of two amplicons from Vinjevatn within an OTU that comprised a total of 279 amplicons. In the second case, the value was found for two amplicons from lake Milsjön in an OTU that comprised a total of 160 amplicons. When many amplicons from the same lake occurred within one OTU, high or low assortativity values where scarcely observed (Figure 4). For example, 618 amplicons from Vostervatnet were placed in OTU #1, but their assortativity within this OTU was only -0.0779. In general, there was no strong indication of biogeographic grouping based on assortativity within any of the 82 Swarm OTUs in this study. For each lake in each individual OTU we ran Kolmogorov-Smirnov tests on the significance of observed assortativity values. For 4130 out of 4369 combinations (89.03%) of OTUs and lakes, assortativity values were significantly different compared to assortativity values obtained from 1,000 randomized structures of the same network (using 95% confidence intervals). In 93.66% of the 4130 cases, the observed assortativity was lower than the 95% confidence interval of the randomized assortativity values; in 6.34% of the 4130 cases, the observed assortativity was higher than the 95% confidence interval of the randomized assortativity values.

**FIGURE 3.**
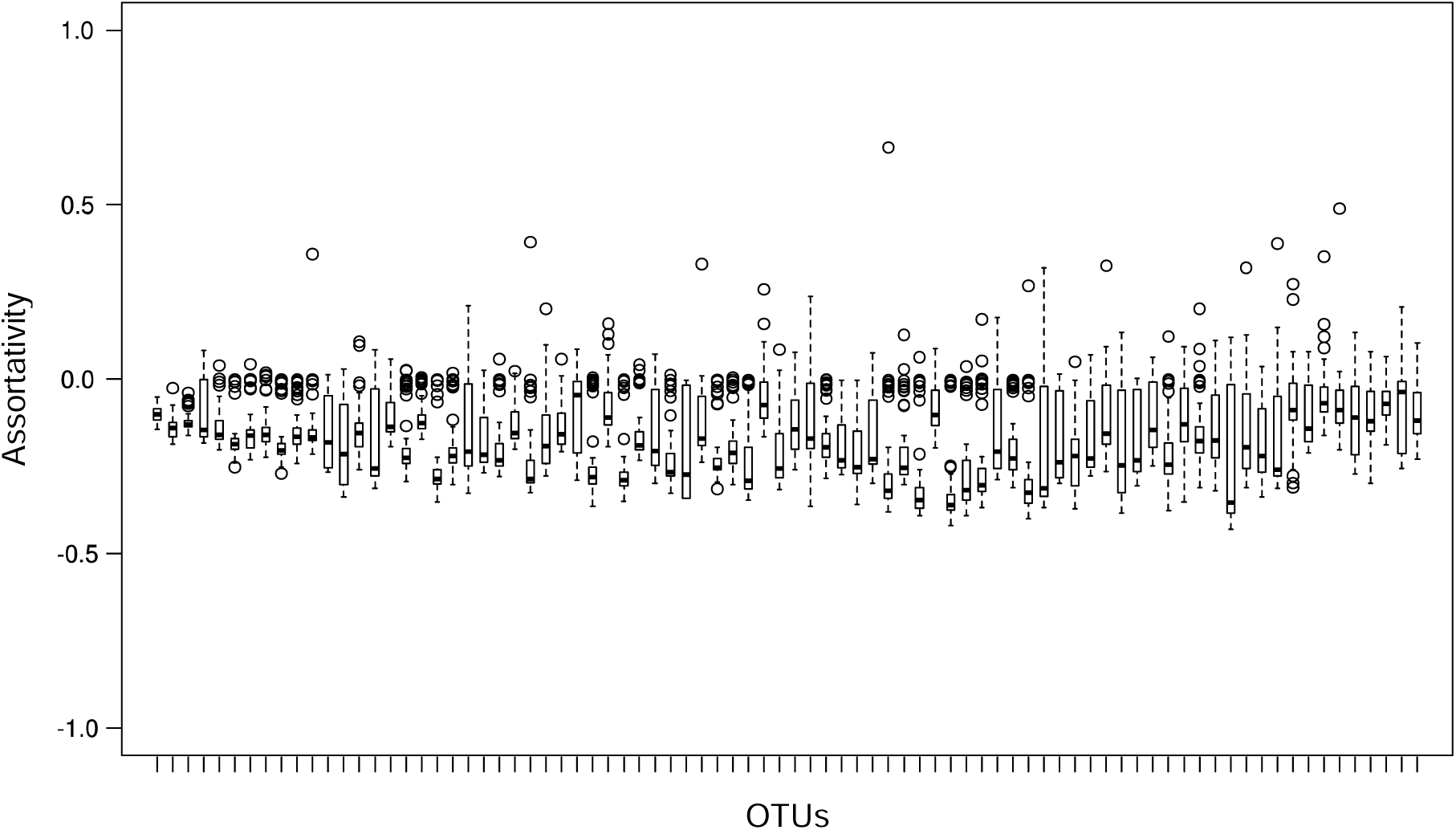
Assortativity of amplicons in individual OTUs. Calculated independently for each lake within each OTU. That is, one boxplot comprises the assortativity values of all lakes obtained from a single OTU.

**FIGURE 4.**
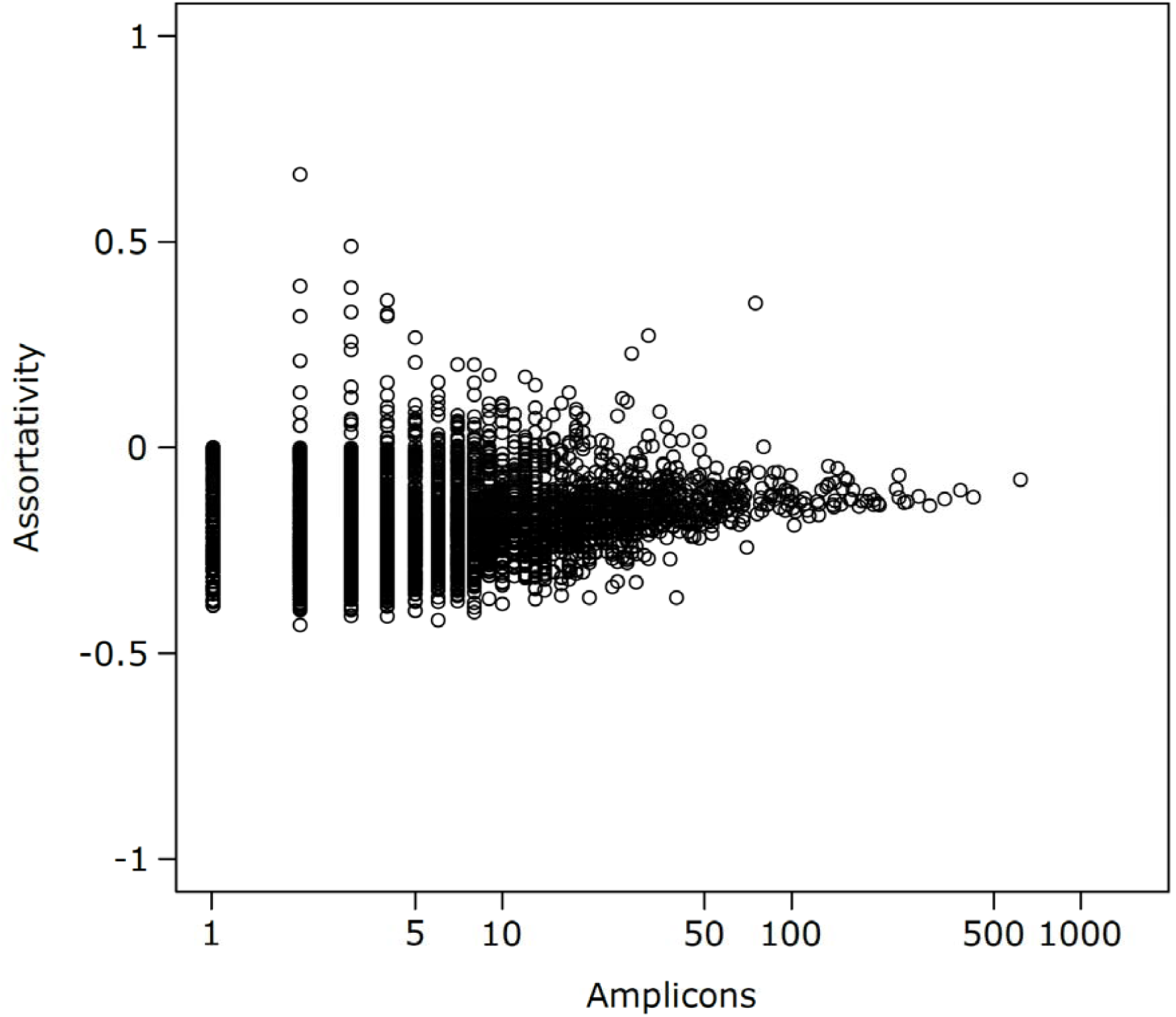
Assortativity of amplicons in relation to their abundance within each OTU. The plot shows the assortativity of each lake’s amplicons within each OTU. Each data point represents the assortativity of a group of amplicons from the same lake within the same OTU. Assortativity values tend towards a value of 0 (indicating random mixing of nodes) when larger groups of amplicons are investigated.

### Geographic assortativity

All widespread OTUs all had their geographic centroid placed at the middle of the study sampling area (i.e., within a 50 km radius around the study centroid), except for OTU #83 with a geographic centroid deviating by 74 km toward east (Figure S4). Geographic distances between amplicon centroids ranged from 0 to 736 km with an average distance within each OTU network at 178 km (± 27 km s.d.). In randomized networks, the same values were recovered (mean: 180 km ± 28 km s.d.). For one single unidentified Dinophyceae OTU (OTU #103, Figure S5), the average distance between neighbor amplicons was significantly lower in the observed compared to randomized networks (Mann-Whitney test, corrected p-value = 1.8 e-4 ± 2.6 e-4 s.d.). For this OTU, the average geographic distance between neighbor amplicon’s centroid was 157 ± 105 km in the observed and 187 ± 112 km in the randomized networks. Notably, this OTU is among the three widespread OTUs with more than 90 % of the amplicons occurring in only one lake (610 out of 669) and among the three widespread OTUs with more than 86% singleton amplicons (579 out of 669), which supports the significant effect is more driven by low abundant amplicons with potential artefactual origin more than by a real biogeographic structure.

## DISCUSSION

### Network analyses can evaluate within-OTU biogeographic variation

Using shortest path analyses, we first asked if amplicons from the same lake are connected directly to each other within the OTUs. Our results showed that the majority of amplicons in the metabarcoding dataset from the Norwegian and Swedish lakes, had indeed at least one neighboring amplicon from the same lake to which they were directly linked in the Swarm OTU networks. Analyses of confidence intervals confirmed some biogeographic structuring of the data, since the mean shortest path distance between two amplicons from the same lake was smaller than expected by chance. Despite this result, shortest path analyses here still emerged as only an indirect measure of biogeographic structuring within the OTUs. Though the results indicate that most amplicons of the same lake occurred at least in pairs (i.e. having a shortest path distance of 1), they do not allow for concluding if these pairs were arranged sequentially, e.g. such as in a chain or even in a clique (such as in Figure 1C). In addition, with shortest path analyses alone, it is unclear if paths of a distance of 1 emerged from scattered pairs or small groups of amplicons from the same lake across many OTUs. Alternatively, they may still indicate biogeographically isolated populations within few OTUs.

Using assortativity analyses, we then asked if amplicons from the same lake form assortative clusters within the OTUs. Assortativity analyses for the complete Swarm OTU network (one value across all OTUs) and for individual OTUs (one value for each OTU), did not show clear assortative or disassortative grouping of amplicons from the same lake into genetically distinct populations. Nevertheless, assortativity analyses emerged here as a more powerful method to address biogeographic variation within OTUs than shortest path analyses. Previous work by Forster et al. (2015) had already demonstrated how assortativity can be used to investigate spatial distribution patterns between OTUs in sequence similarity networks of high-throughput sequencing data. This knowledge was transferred here to a within-OTU application. The analysis delivers a distinct value for inferring whether or not an underlying structure of biogeographic variation is present in a network. Here, assortativity analyses for the complete Swarm OTU network and for individual OTUs, did not show clear assortative or disassortative grouping of amplicons from the same lake into genetically distinct populations. The assortativity of each lake’s amplicons across all OTUs obtained in this study were lower than those obtained by Forster et al. (2015), who found assortativity values as high as 0.7284 when analyzing geographic variation between protistan communities along the European coastline. Fondi et al. (2016) also used assortativity analyses to conclude that ecological niche affiliation was an underlying factor that determined horizontal gene transfer between different types of habitats. The assortativity values observed in Fondi et al. (2016) on a global scale were also higher than those observed by our study on Norwegian and Swedish lakes. Both mentioned studies interpreted the assortative sequence groupings in their dataset as a result of adaptive evolutionary processes caused by environmental factors (habitat and niche affiliation, respectively). If assortative groupings of amplicons by geographic regions could be observed within an OTU, these observations may also be strong indications of evolutionary processes. One Swarm OTU resembles very fine-scaled genetic variation, even on an intraspecific level (D. Forster et al., 2019). Our strategy aimed at testing if this fine-scaled genetic variation is non-randomly distributed but structured by geographic origin. Transferred to an evolutionary framework, one may detect allopatric speciation processes in which intraspecific subpopulations in different geographic regions are characterized by distinct haplotypes. For this, however, time must be sufficient to allow for new mutations from a distinct amplicon to act and produce new neighbors. Using geographic analyses, we asked if amplicons are connected with other amplicons occurring in neighboring lakes. Although this broadened the idea of assortative grouping of amplicons we still observed a lack of biogeographic structuring within OTUs when we used geographic centroids as a proxy to the area covered by an amplicon and by using the distances between those centroids as measure of spatial distribution overlap between amplicons. One single OTU had significant biogeographic structure using this method, but this finding was most likely biased by a high amount of singleton amplicons (i.e., with an abundance of one/observed only once in one lake) linked to each other.

Even though few assortative values were obtained for some OTUs, these do not contradict the lack of biogeographic variation in the dataset. Instead the values corroborate and expand the results of the shortest path analyses: there may be scattered pairs or small groups of amplicons from the same lake that are linked within some OTUs, but there are no larger populations that dominated specific areas within any OTU. From the results of all types of network analyses we conclude that biogeographic distribution is not an underlying structuring factor of genetic diversity within the networks of the most abundant and widespread protist OTUs in lakes of southern Norway and Sweden.

The lack of biogeographic variation recovered by the network analyses approach does not mean that there was generally no biogeographic variation in the dataset. Khomich et al. (Khomich et al., 2017) had previously shown a longitudinal diversity gradient between the OTUs, similar to observations of dispersal patterns in other metabarcoding studies (Fonseca et al., 2014; Lentendu et al., 2018; Ritter et al., 2019; Tedersoo et al., 2014; Venter et al., 2017). Though we achieved an increase of resolution from community level to within-OTU level, the biogeographic variation found among the communities could not be retraced within the network structure of the most widespread OTUs. This suggests that genetic variation within these OTUs does not mirror OTU-specific populations among separated lakes of the same region and, expanding this thought, that there is no indication of allopatric speciation. This also suggest that Swarm’s clustering approach produces ecologically and biologically consistent OTUs, with hypothetical, biogeographically segregated, amplicons from the same species which have accumulate more than one nucleotide difference being efficiently separated into different OTUs. Consequently, the lack of within OTU biogeography cannot readily be extrapolated to OTUs or amplicon sequence variants (ASVs) produced by different approaches (e.g., global clustering algorithm, error-model based grouping algorithm) and remains to be tested.

Some genetic variation within the OTUs may result from intraspecific polymorphism in the hypervariable V4 gene. For some microbial eukaryotes, polymorphisms are even found among the rRNA copies of the same individual (Decelle, Romac, Sasaki, Not, & Mahé, 2014; Gong, Dong, Liu, & Massana, 2013; Weber & Pawlowski, 2014). It is further possible that genetic variation within the OTUs results from gene flow between the lake habitats. Microbial organisms are known to migrate with the help of other organisms. Fish, birds and humans are some of the potential vector organisms that maintain gene flow between lakes by mediating the migration of microbial species (Coughlan, Kelly, Davenport, & Jansen, 2017; Foissner, 2011; Kristiansen, 1996). Most likely, the scale on which we evaluated biogeographic variation was not large enough for contrasting genetically different populations within the most widespread OTUs (if there are any). Intraspecific diversity studies of microbial eukaryotes on a global scale could successfully distinguish between genetically different populations and show that there are limits to gene flow that lead to biogeographically distinct genetic variants of the same organism (Casteleyn et al., 2010; Kim, Park, Kim, Wang, & Han, 2015). In bacteria, some widespread generalist species were identified as ancestral lineages from which several specialist species descended after spreading into new environments (Sriswasdi, Yang, & Iwasaki, 2017). The results of our network analyses did not confirm a similar adaptive radiation for microbial eukaryotes in Norwegian and Swedish lakes.

Network-based analyses of intraspecific variation have a long tradition in phylogeography and population genetics. In these previous approaches, an alignment or a distance matrix is used as the basis for phylogenetic network inference (Huson & Bryant, 2006). The partitioning of genetic variation within the network is then used to explore patterns of geographical association as well as the frequency of central vs. low-frequency derived haplotypes to address demographic questions such as population expansions and bottlenecks (Raupach et al., 2010). Besides their utility to explore genetic data, phylogenetic networks are also frequently used to systematically partition intraspecific variation in the network using a Nested-Clade Analysis (NCA) or Nested-Clade Phylogeographic Analysis (NCPA) approach (Templeton, 1998; Templeton, Routman, & Phillips, 1995). In the NCA approach, the network is sub-structured into hierarchically nested sub-networks (clades) inside a parsimony network. The aim of NCA is to test the probability of different microevolutionary processes (e.g., isolation in allopatry, restricted gene flow, long-distance dispersal, panmixia). The test treats sample locations as categorical variables and estimates the probability of association using an exact permutational contingency test. In addition, the geographic distance between the locations can be used for the test statistics to assess the likelihood of long-distance dispersal and other processes (Posada, Crandall, & Templeton, 2006). While NCA is criticized for having high error rates when using simulated data, the proposed expectations of the NCA to test for are still valid. The approach proposed here differs from NCA in that it does not perform specific nestings, but rather uses assortativity; it only uses presence-absence assessment per population rather than the integration of all networks into one; and it integrates geographical patterns of within-OTU variation for a metabarcoding set consisting of up to thousands of data points.

### Wider applicability of within-OTU network analyses

While the application of assortativity analyses for inferring biogeographic variation in metabarcoding datasets has been successfully demonstrated before (Fondi et al., 2016; D. Forster et al., 2015), our workflow is the first combination of assortativity analyses within-OTU networks created by Swarm. Our application is also the first to exploit Swarm’s feature of fine-grained single-nucleotide differences for clustering instead of comparably coarse-grained clustering via fixed sequence similarity thresholds. The test for biogeographic variation presented here is a statistical expansion to evaluate Swarm clustering results.

Commands for reproducing this step are available at https://github.com/lentendu/DeltaMP and do not demand additional pre- or post-clustering data treatment. Although we specifically chose Swarm because its OTU structure was well-suited for network analyses, the application of assortativity and shortest path analyses is adaptable to the results of other clustering algorithms. Ideally, the results of these algorithms should also exhibit an underlying network structure or provide information from which a network may be inferred. For instance, the results of two widely used and powerful clustering algorithms are not directly suited for within-OTU network analyses. USEARCH (Edgar, 2010) relies on heuristic clustering and lacks information for all pairs of sequences. DADA2 (Callahan et al., 2016), which is becoming more and more popular in metabarcoding studies, uses strict error-correction models for creating amplicon sequence variants (ASVs), but provides little information for evaluating the genetic diversity within the ASVs. For these two examples, an additional step for obtaining network structures would have to be implemented (D. Forster et al., 2019), before network analyses may be performed.

## ACKNOWLEDGEMENTS

We thank Eric Bapteste for helpful comments on an earlier draft of the manuscript. Funding information: Carl Zeiss Foundation to DF, and Deutsche Forschungsgemeinschaft grant DU1319/5-1 to MD.

## DATA ACCESSIBILITY

Codes for bioinformatic pipeline is available at https://github.com/lentendu/DeltaMP (v0.4). The R script used to perform statistical analyses and create the figures (File S1) and the configuration file used to run the bioinformatic pipeline (File S2) are provided as Supporting information in text format. The full OTU table produced by the bioinformatic pipeline and used for the statistical analyses (Dataset S1) as well as Swarm’s OTU internal structure (Dataset S2), Swarm’s OTU amplicon network file (Dataset S3) and the amplicon sequence table (Dataset S4) are provided as supplements. Raw nucleotide sequences with corresponding mapping files, originally from Khomich et al. (2017), are available at Dryad (doi:10.5061/dryad.7s6s8).

## Supplementary Table & Figures

**Table S1** Summary of shortest path analyses for amplicons of each lake across all OTUs.

**FIGURE S1.**
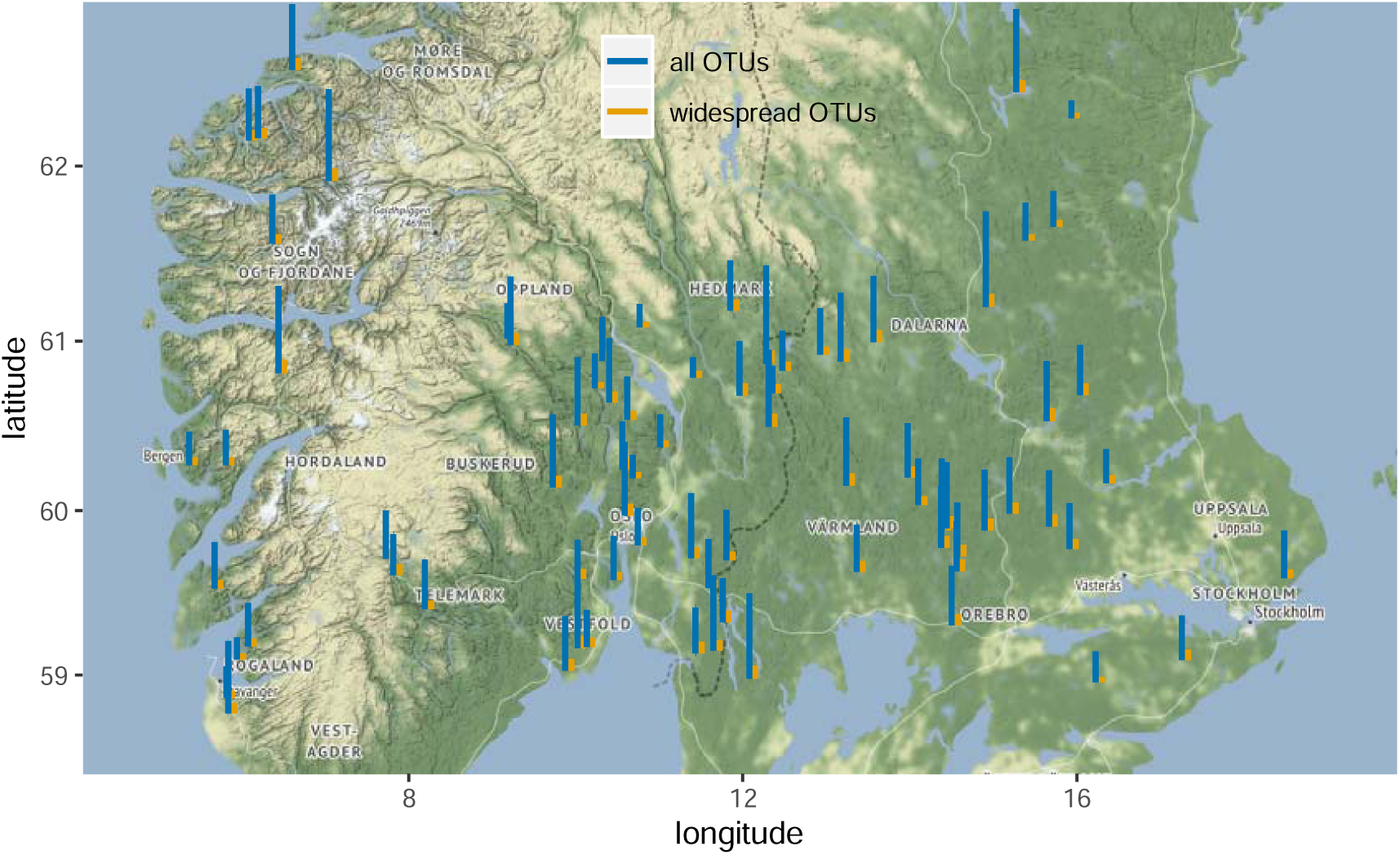
Map of the sampled lakes with barplots of the OTU richness placed just above the geographic coordinates of each lake. OTU richness bars were log-transformed to better visualize both types of OTU richness: OTU richness of all protists ranged from 98 to 571 (blue bars); OTU richness of widespread protists ranged from 31 to 80 (orange bars).

**FIGURE S2.**
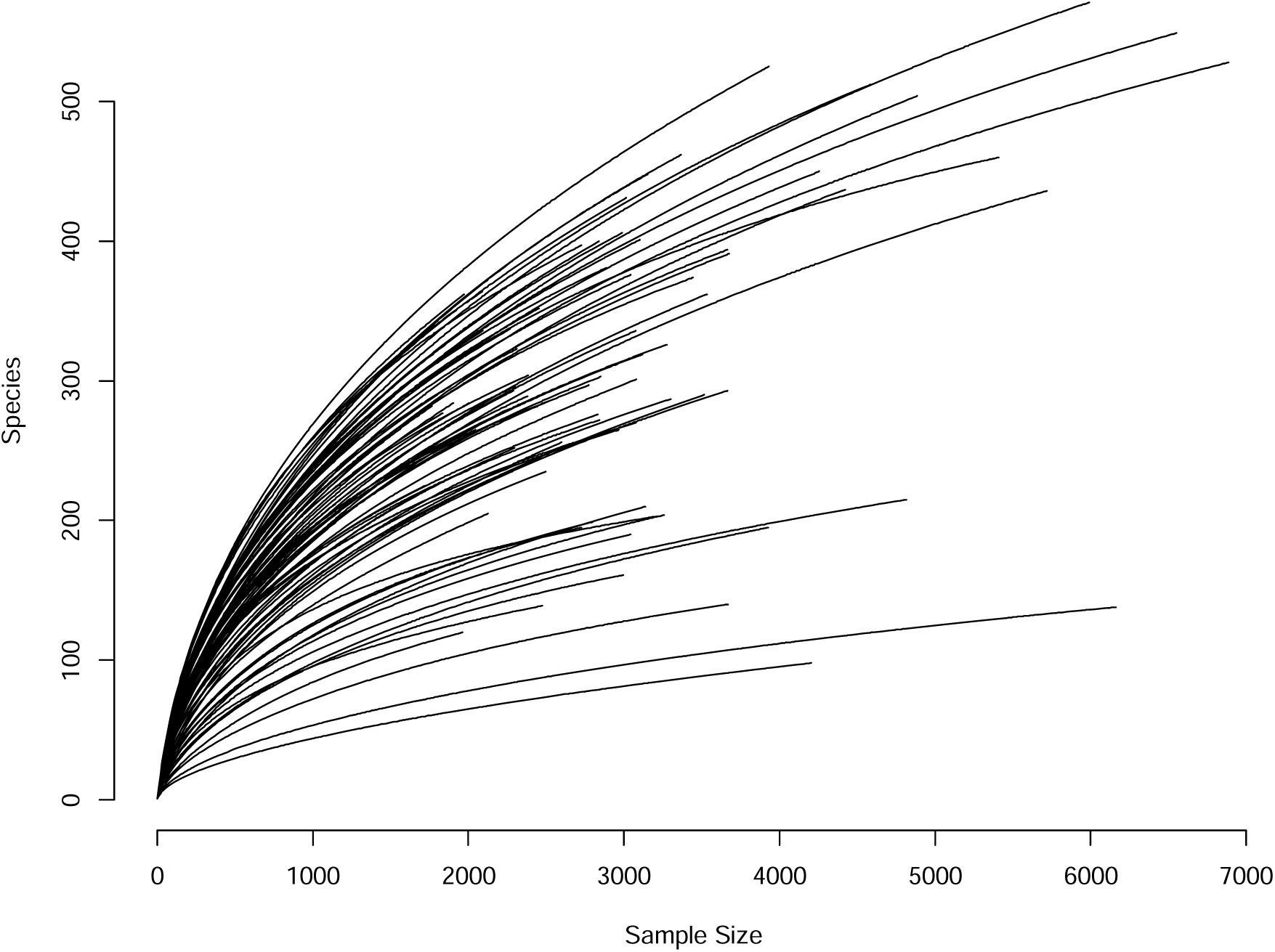
Rarefaction curves of all samples in the dataset.

**FIGURE S3.**
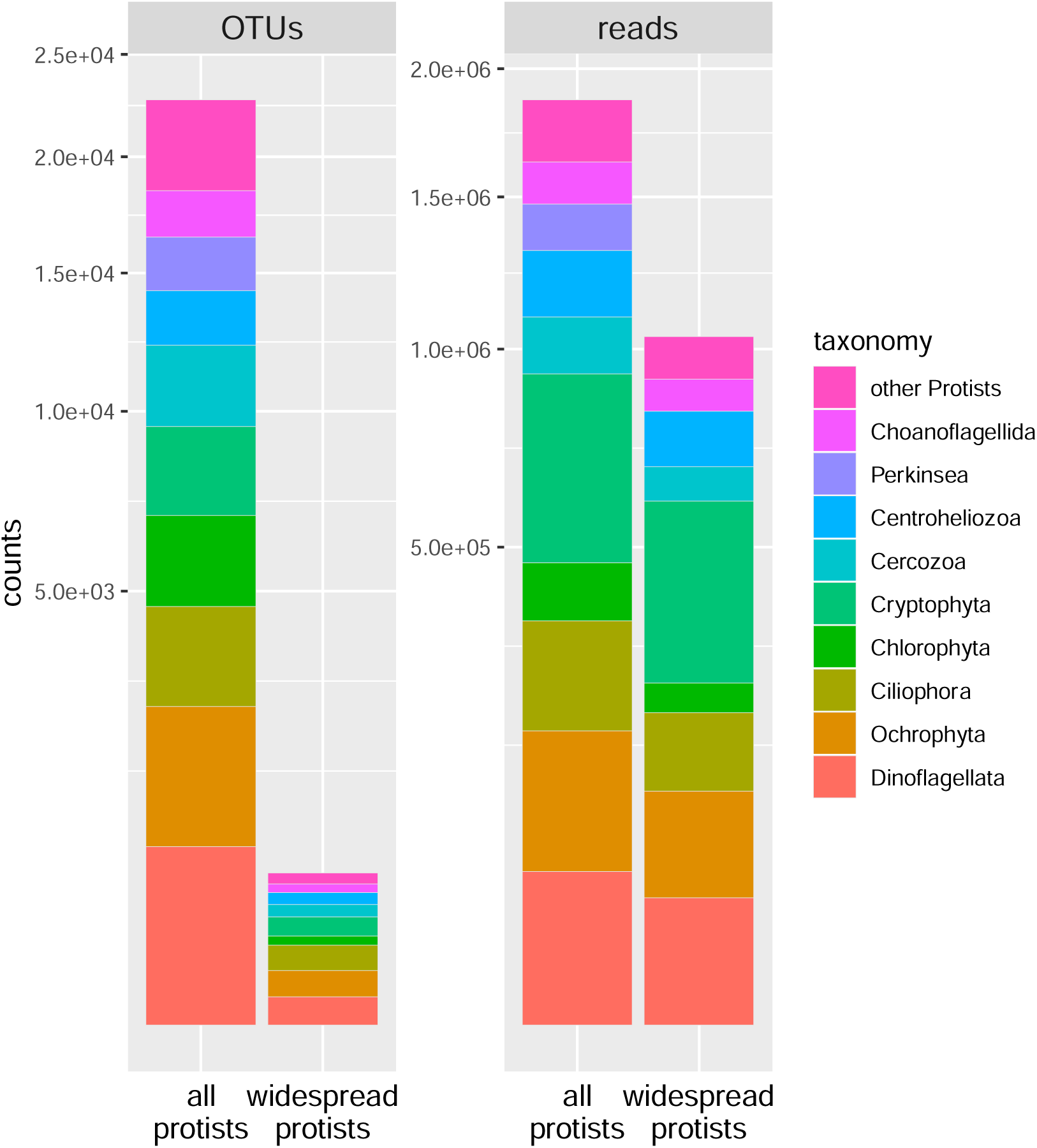
Taxonomic composition of all protist OTUs and widespread protist OTUs in term of OTU and read counts. Y-axis scale was square-root transformed to ease visualization of counts for the widespread protistan OTUs.

**FIGURE S4.**
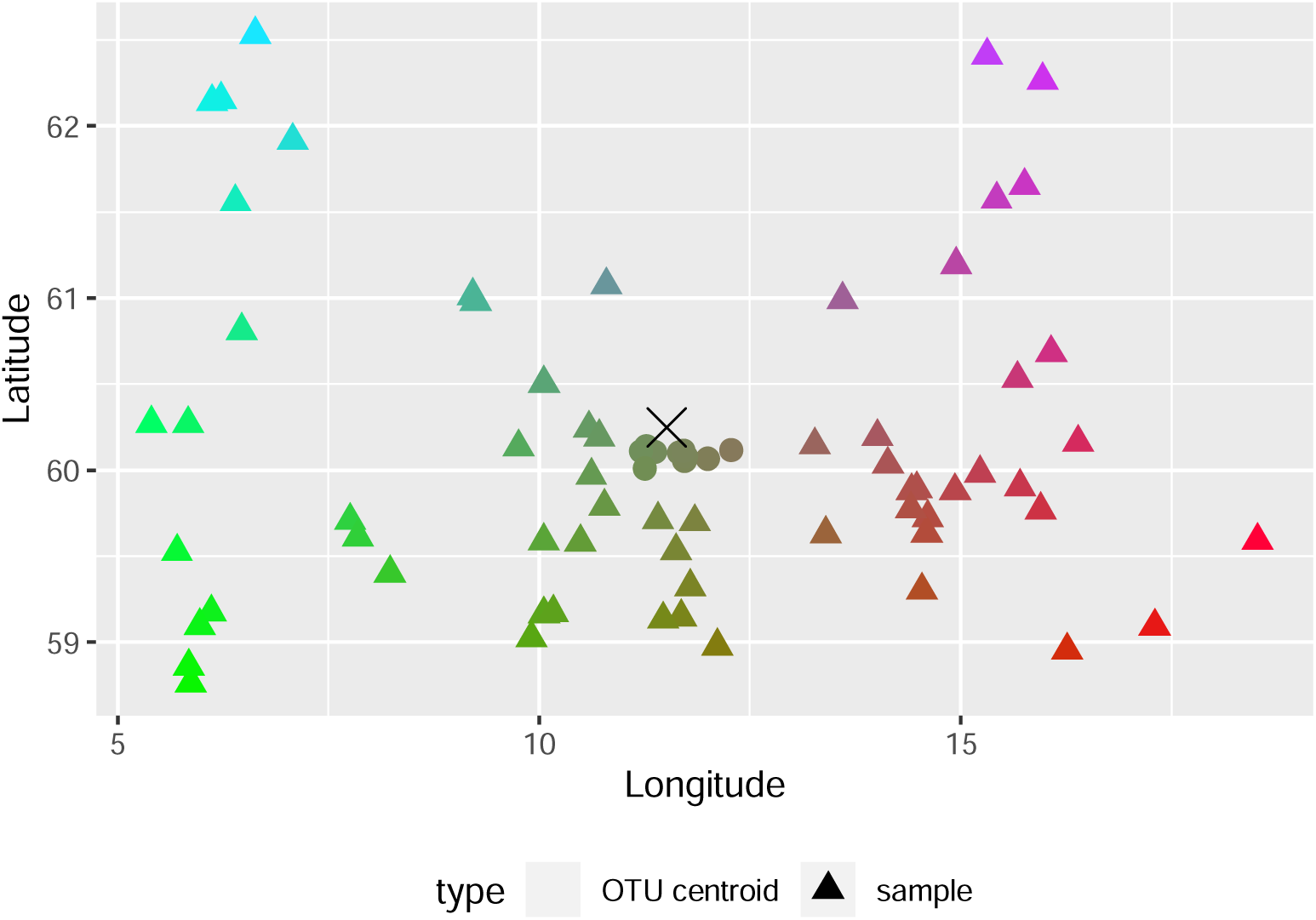
Sample geographic positions and widespread protistan OTU centroid geographic position. OTU centroid is the centroid geographic location of all lakes an OTU is occurring in. The color scale is a 2-dimensional gradient spanning the sampling area (see Figure S5).

**FIGURE S5.**
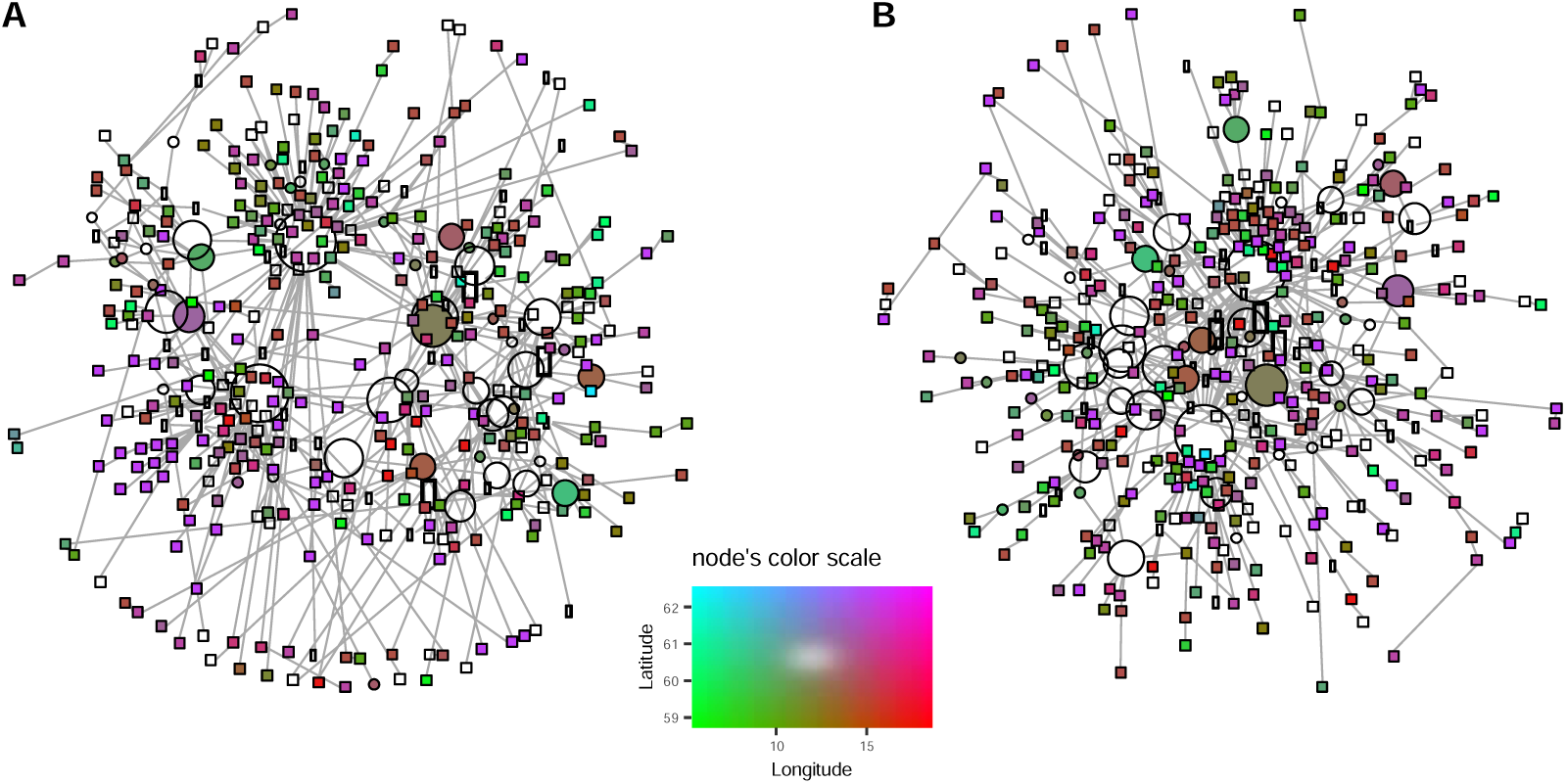
Sequence similarity network of OTU #103 using observed edges (A) and one example of randomized edges (B). Nodes represent singleton (square) and non-singleton (circle) amplicons. Node colors are adapted from the 2-dimensional color gradient of the legend using their centroid geographic location. Edges represent a single nucleotide difference between linked amplicons. The edge length is proportional to the distance between geographic centroids of connected amplicons.

